# A deep learning model embedded framework to distinguish DNA and RNA mutations directly from RNA-seq

**DOI:** 10.1101/2023.10.17.562625

**Authors:** Zhi-Can Fu, Bao-Qing Gao, Fang Nan, Xu-Kai Ma, Li Yang

**Author notes:** These authors contributed equally to this work (Zhi-Can Fu, Bao-Qing Gao).

## Abstract

We develop a stepwise computational framework, called DEMINING, to directly <u>d</u>etect <u>e</u>xpressed DNA and RNA <u>m</u>utations in R<u>N</u>A deep sequenc<u>ing</u> data. DEMINING incorporates a deep learning model named DeepDDR, which facilitates the separation of expressed DNA mutations from RNA mutations after RNA-seq read mapping and pileup. When applied in RNA-seq of acute myeloid leukemia patients, DEMINING uncovered previously-underappreciated DNA and RNA mutations, some associated with the upregulated expression of host genes or the production of neoantigens. Finally, we demonstrate that DEMINING could precisely classify DNA and RNA mutations in RNA-seq data from non-primate species through the utilization of transfer learning.

## Introduction

A myriad of DNA and RNA variants that are different to the human reference genome have been identified by analyzing fast-accumulated high-throughput sequencing datasets^1, 2^, referred to as DNA mutations (DMs) and RNA mutations (RMs), respectively. Other than directly called from high-quality whole genome sequencing (WGS) datasets^1^, DMs could be also interpreted in RNA-seq datasets^3, 4^, but were generally treated as noises during profiling RMs, most of which are Adenosine-to-Inosine (A-to-I) RNA editing catalyzed by adenosine deaminase acting on RNA (ADAR) enzymes^5^. Indeed, calls in A-to-I RMs from RNA-seq datasets are fraught with difficulties of expressed DMs and other artifacts including sequencing and mapping errors^6^. To solve these problems, stepwise bioinformatic tools have been developed to precisely identify A-to-I RMs from RNA-seq datasets by using parallel WGS and/or prior-knowledge for the removal of DMs^7^. Moreover, additional tools have been developed for A-to-I RMs calling without the requirement of matched WGS, achieved by analyzing multiple RNA-seq datasets for common/constitutive A-to-I RMs or taking advantage of variable allelic linkage between DMs and A-to-I RMs from reads mapped to the same regions^8, 9, 10^.

Given the fact that RNA is transcribed from DNA, it is luring to identify disease-associated genetic DMs directly from RNA-seq datasets. To achieve this goal, we developed a stepwise computational framework to directly <u>d</u>etect <u>e</u>xpressed DNA and RNA <u>m</u>utations in R<u>N</u>A deep sequenc<u>ing</u> datasets, referred to as DEMINING. DEMINING incorporated a deep learning model named DeepDDR, which achieved the differentiation of expressed DMs from RMs directly from aligned RNA-seq reads. When applied in RNA-seq of acute myeloid leukemia (AML) patients, DEMINING uncovered previously-underappreciated DMs and RMs in unannotated AML-associated gene loci. The association of these DMs (and RMs) with their host gene upregulation and the production of neoantigens suggested possible targets for diagnosis and therapeutics of the AML disease. Finally, after transfer learning, DeepDDR-transfer embedded DEMINING achieved successful classification of DMs and RMs from aligned RNA-seq reads in other species, such as mouse and nematode.

## Results

### Development of DEMINING to differenti te DMs and RMs directly from RNA-seq datasets with the embedded DeepDDR model

To identify expressed DMs and RMs directly from RNA-seq datasets, DEMINING first called high-confidence mutation candidates with RNA-seq read mapping and filtering, and then fed them into the featured DeepDDR model for DM and RM classification (Fig. 1a, Methods). The DeepDDR model was trained with reliable sets of DMs and RMs obtained from paired human WGS and RNA-seq datasets of 403 donors (Supplementary Table 1), retrieved from the 1000 Genomes Project^1^ and the Geuvadis consortium^11^, respectively. Of note, among ∼ 31 millions of high-confidence DMs provided by 1000 Genomes Project, about 578 thousands of them could be examined in the paired 403 RNA-seq datasets with both expression (≥ 3 hits per billion mapped bases^10^, HPB) and mutation frequency (≥ 0.05 cutoffs, and similar distributions of all twelve mutation types were observed between samples (Supplementary Fig. 1a, Methods). High-confidence RMs in each of 403 RNA-seq datasets were individually detected by the previously-published RADAR pipeline^12^, and further filtered by paired WGS dataset to remove DMs (Supplementary Fig. 1b, Methods). The same expression (≥ 3 HPB) and mutation frequency (≥ 0.05) filters were used for the selection of high-confidence RMs. Of note, over 98% of identified RMs in each of 403 RNA-seq datasets were A-to-G and T-to-C mismatches, mainly in *Alu* regions, indicating high-confidence A-to-I RMs^7^ called by RADAR^12^ (Supplementary Fig. 1b). High-confidence A-to-I RMs identified by the RADAR pipeline in each of 403 RNA-seq datasets were then combined together to generate the collection of 122,872 unique RMs (Supplementary Table 2). In parallel, 122,872 unique DMs were randomly selected from 578 thousands of expressed DMs (Supplementary Table 3). As expected, mutation frequencies of DMs were enriched at 1 or 0.5, while those of RMs were largely below 0.5 (Supplementary Fig. 1c). These DMs and RMs were then individually split into training, validation, and test sets with an 8:1:1 ratio for DeepDDR development and evaluation (Supplementary Fig. 1d).

**Fig. 1.**
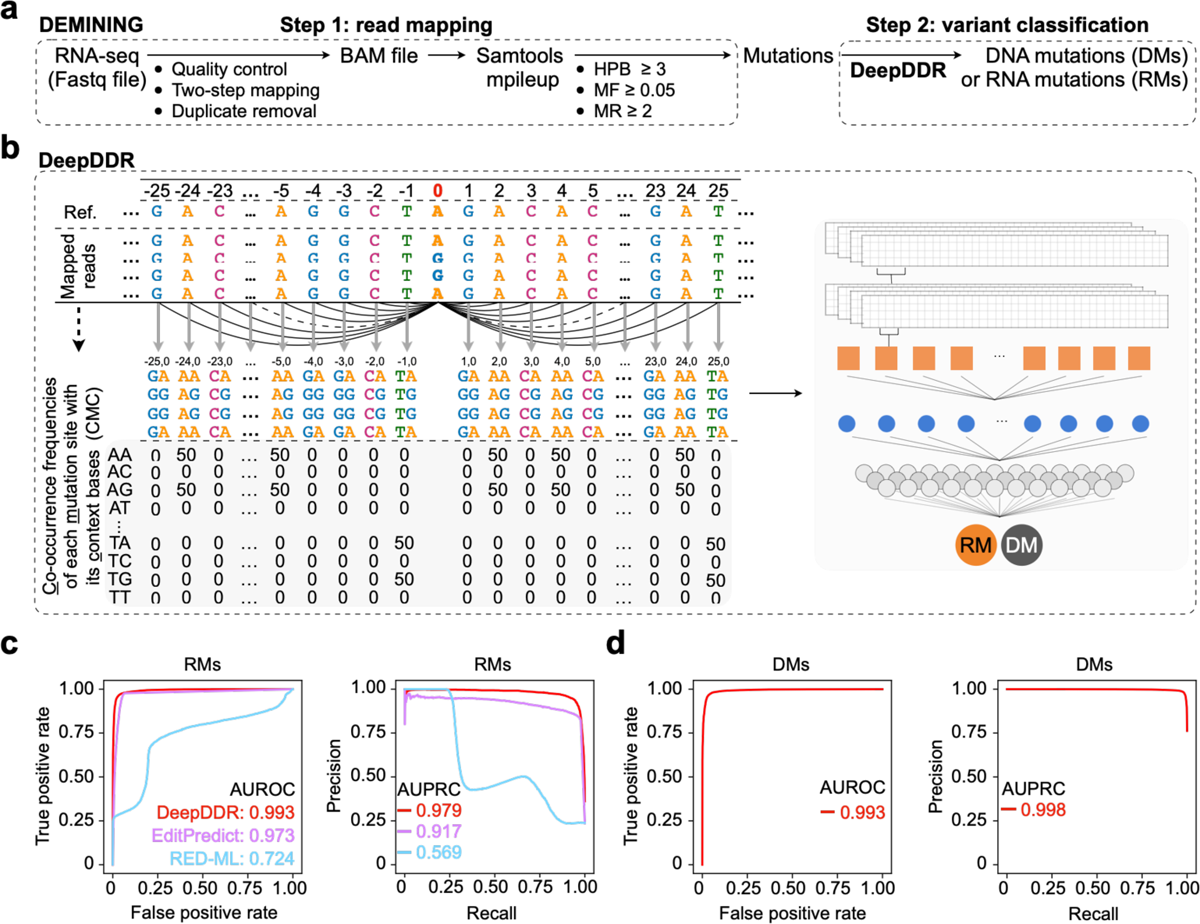
Developing DEMINING embedded DeepDDR model for DNA mutations (DMs) and RNA mutations (RMs) classification. **a**, Construction of a stepwise DEMINING computational framework for direct DNA mutation (DM) and RNA mutation (RM) classification. HPB: hits per billion mapped bases, MF: mutation frequency, MR: mutation read. See “Methods” section for details. **b**, Schematic diagram of an embedded DeepDDR model for DM and RM classification. Left, features extract strategy by the <u>c</u>o-occurrence frequencies of each <u>m</u>utation site with its <u>c</u>ontext bases (CMC). Right, DeepDDR model architecture. See “Methods” section for details. **c**, Evaluation of different models on RM identification. Receiver Operating Characteristic (ROC, left) curves and Precision Recall Curves (PRC, right) of DeepDDR (red), EditPredict (purple) and RED-ML (blue) were shown to indicate their performance on RM identification with an independent sample (HG00145). Area Under ROC (AUROC) and Area Under PRC (AUPRC) values of DeepDDR (red), EditPredict (purple) and RED-ML (blue) were included in the figure. **d**, Evaluation of DeepDDR on DM identification. ROC (left) and PRC (right) of DeepDDR were shown to indicate its performance on DM identification with the same independent sample (HG00145). AUROC and AUPRC values of DeepDDR were included in the figure.

To extract features of these mutations, 51-nucleotide sequences containing a given DM (or RM) in the middle and its 50 context bases (25 bases in the upstream region and 25 bases in the downstream region) were first piled up from reads mapped to the human reference genome (hg38) in aligned BAM files (Fig. 1b). Next, a matrix of the <u>c</u>o-occurrence frequencies of each <u>m</u>utation site with its <u>c</u>ontext bases (CMC) was calculated, and CMC matrices of all DMs and RMs in the training set were used as the input to train the DeepDDR model that contains two layers of convolutional neural network (CNN) (Fig. 1b, Methods).

DeepDDR outputs a prediction probability from 0 to 1 for a given DM or RM, followed by the classification into the DM group (with prediction probabilities ≥ 0.5) or the RM group (with prediction probabilities < 0.5). It was worthwhile noting, the majority (> 98%) of DMs were shown with prediction probabilities between 0.5 to 1 and most (> 92%) RMs with prediction probabilities between 0 to 0.5 in the test set (Supplementary Fig. 2a). Compared to previously-published EditPredict and RED-ML methods that were designed only for A-to-I RM prediction, DeepDDR could find more true RMs as well (Supplementary Fig. 2a-c). Meanwhile, DeepDDR achieved AUROC (Area Under the Receiver Operating Characteristic curve) values over 0.99 on both RM (Supplementary Fig. 2d) and DM (Supplementary Fig. 2e) prediction, much higher than previously-reported EditPredict (https://github.com/wjd198605/EditPredict) and RED-ML (https://github.com/BGIRED/RED-ML) methods that only achieve RM prediction.

In addition, an independent test set (SampleID: HG00145 from the 1000 Genomes Project and the Geuvadis consortium) was downloaded for additional evaluation. With the same pipelines (Supplementary Fig. 1a and 1b), true DMs and RMs were individually extracted from this paired WGS and RNA-seq datasets, and then used to evaluate predicted DMs/RMs by DEMINING from the RNA-seq only (Supplementary Table 4). Similar to results evaluated by the test set, DeepDDR outperformed EditPredict and RED-ML, with the highest AUROC and AUPRC values on the prediction of RMs (Fig. 1c, Supplementary Fig. 3a-d). In addition, DeepDDR performed equally well in terms of AUROC and AUPRC values for predicting DMs (Fig. 1d and Supplementary Fig. 3e). Of note, different to EditPredict and RED-ML designed to call (A-to-I) RMs only, DeepDDR could directly predict and distinguish DMs and RMs from mapped RNA-seq reads, suggesting the uniqueness and superiority of using DeepDDR for efficient DM and RM classification from publicly-available RNA-seq datasets.

### Application of DEMINING to detect AML-associated DMs/RMs from patient samples

Next, we set to apply the DeepDDR-embedded DEMINING framework to identify disease-related mutations from publicly available RNA-seq datasets of individuals, more specifically, AML patient samples in this study. From a published collection of 19 AML patient RNA-seq datasets^13^ (peripheral blood or bone marrow tissue), DEMINING totally identified 195,256 DMs (Fig. 2a and Supplementary Table 5) and 137,682 RMs (Supplementary Table 6). The heterogeneous nature of AML has been associated with numerous pathogenesis-related mutations^14^, prompting us to further investigate these DEMINING-identified mutations from AML samples.

**Fig. 2.**
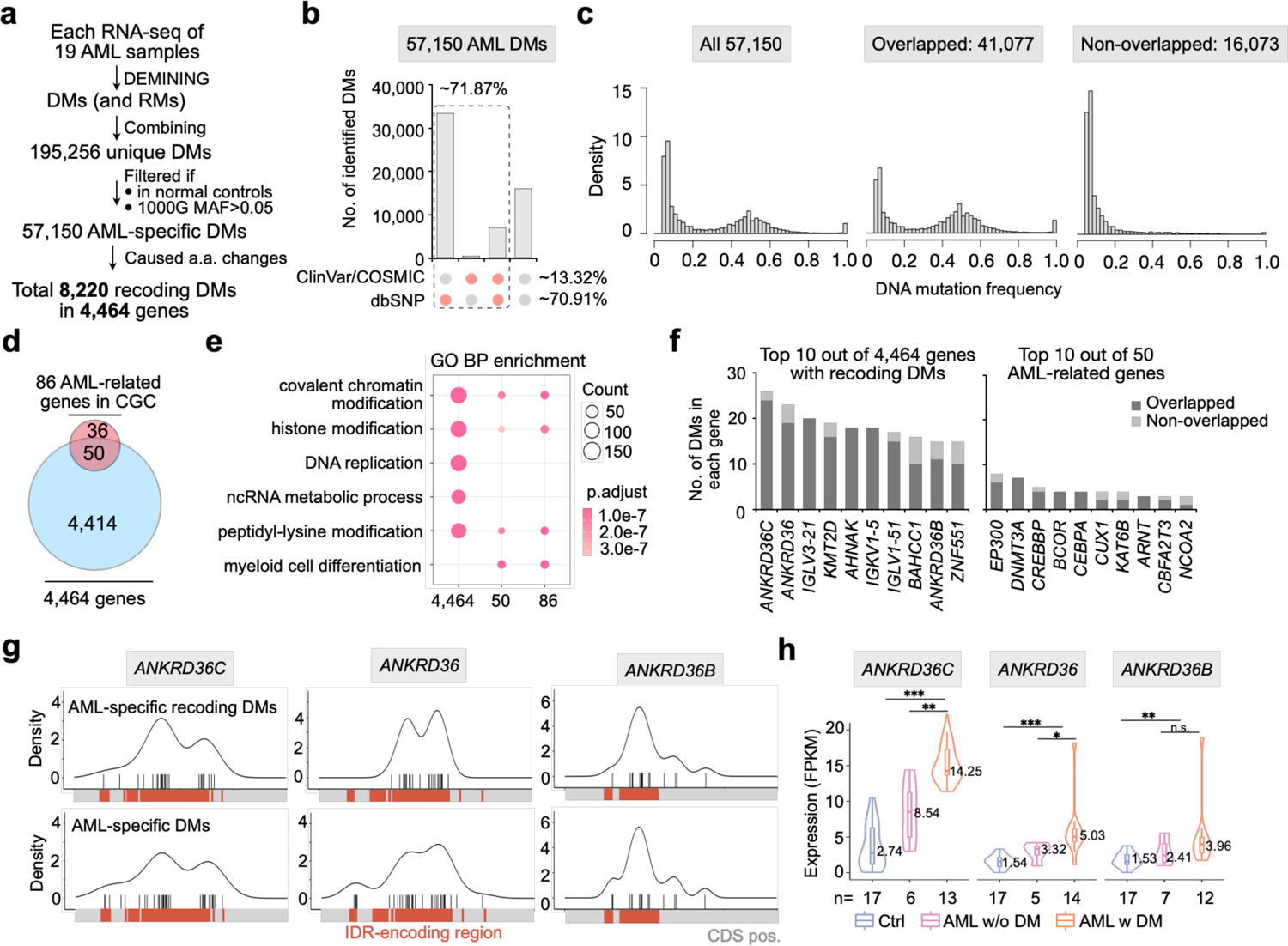
Applying DEMINING framework to identify disease-related mutations in acute myeloid leukemia (AML). **a**, Identification of AML-associated DMs from corresponding RNA-seq datasets^13^ by DEMINING. **b**, Overlapping of AML-specific DMs and reported SNVs in public databases, including ClinVar 202304 (https://www.ncbi.nlm.nih.gov/clinvar/), COSMIC (version 97, https://cancer.sanger.ac.uk/cosmic/) and dbSNP (version 156, https://www.ncbi.nlm.nih.gov/snp/). **c**, Mutation frequency distribution of all AML-specific DMs (left), overlapped AML-specific DMs (middle) and non-overlapped AML-specific DMs (right). **d**, Overlapping of 4,464 mutated genes carrying AML-specific recoding DMs with 50 AML-associated genes listed in COSMIC Cancer Gene Consensus (CGC). **e**, Gene Ontology (GO) enrichment analysis in biological process (BP) terms for three gene sets including all mutated 4,464 genes, 86 AML-associated genes listed in CGC, and their overlapping 50 genes. Top GO terms ordered by adjusted *P* value in at least one gene set were kept and compared. **f**, The number of overlapped (dark grey) and non-overlapped (light grey) DMs in the top ten genes. Left, top ten genes out of 4,464 genes with recoding DMs; right, top ten genes out of 50 AML-associated genes listed in COSMIC CGC. **g**, Distribution of identified AML-specific recoding DMs (top) and AML-specific DMs (bottom) along coding sequences (CDS) of the *ANKRD36C* (left), *ANKRD36* (middle) and *ANKRD36B* (right). Distribution (black plot) of predicted DMs (black vertical lines) by DEMINING were shown in genes’ CDS regions (grey and red rectangles). Red rectangles represent CDS regions that encoding intrinsically disordered regions (IDRs) predicted by MobiDB-lite integrated in the InterPro database (https://www.ebi.ac.uk/interpro/). **h**, Comparison of gene expression of *ANKRD36C*, *ANKRD36* and *ANKRD36B* in 17 normal control samples (Ctrl) and 19 AML patients with or without recoding DMs (AML w DM: AML patients with recoding DM; AML w/o DM: AML patients without recoding DM). The boxplot summarizes results for all samples with the number of samples n shown below. Center line: Median. Box bottom and top edges: 25th and 75th percentiles. Whiskers extend to extreme points excluding outliers (1.5 times above or below the interquartile range). Outliers omitted for clarity. Violin-shaped areas: Kernel density estimate of data distribution. Statistical significance was assessed with two-tailed Wilcoxon rank-sum test. \**P* < 0.05, \*\**P* < 0.01, \*\*\**P* < 0.001.

Among 195,256 DMs, we first filtered out those could be also called in at least one of 17 normal control samples (medullary thymic or myelocytic precursor cells) in the same collection^13^ or had a high minor allele frequency (MAF > 0.05) in 1000 Genomes Project (Fig. 2a). As a result, 57,150 DMs were highlighted as AML-specific ones (Fig. 2a, Supplementary Table 7), with a similar distribution pattern of mutation types (Supplementary Fig. 4a) as those retrieved from 1000 Genomes Project (Supplementary Fig. 1a). Although the majority (∼70.91%) of them were found to be overlapped with annotated SNPs in dbSNP (Fig. 2b), only a small portion (∼13.32%) of 57,150 AML-specific DMs were previously-reported to be associated with diseases by ClinVar and/or COSMIC databases. These results thus indicated the reliability of using DEMINING to identify DMs directly from RNA-seq, and further implied that many previously-underappreciated DMs might be associated with AML pathogenesis, while their links with AML need further investigation. Nevertheless, mutation frequencies of these previously-unannotated DMs were expectedly lower than those of overlapped ones (Fig. 2c), suggesting they were likely somatic mutations during AML pathogenesis.

Among 57,150 AML-specific DMs identified by DEMINING, 8,220 of them were predicted to cause amino acid changes, hereafter called recoding DMs. These recoding DMs were found in 4,464 genes (Fig. 2d), including 50 out of 86 AML-associated genes listed in the COSMIC Cancer Gene Consensus (CGC) that have evidence-based functions in the development and progression of AML. Importantly, similar biological pathways (BPs) were enriched by Gene Ontology (GO) analyses of 4,464 genes with AML-specific recoding DMs, 86 AML-associated genes listed in CGC, and their overlapping 50 genes (Fig. 2e). These enriched GO pathways included covalent chromatin modification, histone modification and peptidyl−lysine modification (Fig. 2e), which were also found to be mis-regulated during tumorigenesis in previous studies^15^.

### Correlation of DEMINING-identified DMs and mis-regulated gene expression

We further analyzed the distribution of these recoding DMs in their host genes, and observed that over fifteen recoding DMs were enriched in top ten out of aforementioned 4,464 genes (left panel of Fig. 2f, Supplementary Fig. 4b), compared to less than ten recoding DMs in AML-associated genes reported by CGC (right panel of Fig. 2f). Top ten genes with most enriched recoding DMs included *ANKRD36C* (26 recoding DMs), *ANKRD36* (23 recoding DMs) and *ANKRD36B* (15 recoding DMs), which all belong to the ankyrin repeat domain (*ANKRD*) gene family. Interestingly, recoding DMs and all AML-specific DMs identified by DEMINING in the three *ANKRD* gene loci were mainly clustered in predicted sequences encoding intrinsically disordered regions (IDRs) (Fig. 2g). Previous studies showed that some mutations mapped to sequences encoding predicted IDRs were related to diseases^16^. Since IDRs could drive liquid-liquid phase separation^17^ (LLPS) and the dysregulation of LLPS was a key event in the initiation and/or evolution of cancer^18^, the identification of recoding DMs in regions encoding predicted IDRs of *ANKRD* gene loci suggested the interplay of dysregulated LLPS by mutated *ANKRD* genes along AML pathogenesis. Together with a recent finding of *ANKRD36* as a biomarker of Chronic Myeloid Leukemia^19^ (CML), these results strongly suggested the possible association of enriched recoding DMs in *ANKRD* genes with AML.

What was the consequence of these recoding DMs on their host gene expression? We observed all three *ANKRD* genes were significantly up-regulated in AML samples, especially in those with DEMINING-identified recoding DMs (Fig. 2h). These results indicated that DEMINING-identified DMs might be functional as *cis*-expression quantitative trait loci (*cis*-eQTLs), while we could not exclude the possibility of indirect effects of DMs on their host gene expression.

The cluster of 8,220 AML-specific recoding DMs was selected due to their capacity of causing amino acid changes (Fig. 2a). Although lack of the whole-proteome sequencing datasets, corresponding liquid chromatography–coupled mass spectrometry (LC-MS/MS) spectra of major histocompatibility complex (MHC)-associated peptides from 19 AML patients were available^13^. Thus, it was attractive to examine whether 8,220 DEMINING-identified AML-specific recoding DMs (and recoding 149 RMs, Supplementary Fig. 5, Supplementary Tables 7 and 8) could cause the production of neoantigen(s) that were displayed by MHC. To achieve this goal, we created an *in silico* library containing 3,581,675 of predicted peptides (8-30 amino acids (a.a.) long) from 4,464 genes with 8,220 recoding DMs (in Fig. 2a) and 121 genes with 149 recoding RMs (Supplementary Fig. 5) as queries (Supplementary Fig. 6a, Methods). Decoyed peptides from the predicted peptides and proteins from human UniProt proteome were also added to this library to be used as negative controls (Supplementary Fig. 6a, Methods). This *in silico* library containing both predicted peptides with recoding DMs/RMs and negative control peptides was used as a bait to screen LC-MS/MS spectra of MHC-associated peptides from AML patients. This analysis led to the identification of three neoantigen candidates, two caused by distinct DMs and one caused by an RM (Supplementary Fig. 6b). Of note, none of these three neoantigens were listed in the published analysis of LC-MS/MS spectra^13^, highlighting the importance of DM/RM-guided strategy for neoantigen identification.

### Extended application of DEMINING in non-primate samples after transfer learning

Other than the successful application in human samples, we tempted to apply DEMINING to identify DMs and RMs from other non-primate datasets, which was challenging as characteristics of A-to-I RMs in primates, such as human, and non-primates, such as mouse, are very different^20^. Basically, distinct to the large number of human A-to-I and their predominant distribution in primate-specific *Alu* regions, the number of A-to-I in mouse is much smaller, possibly due to the lack of *Alu* elements in the mouse genome^20^. Nevertheless, we adopted DEMINING to differentiate DMs and RMs from a publicly-available mouse bone marrow RNA-seq dataset (Methods, Fig. 3a, top). Because no corresponding WGS was available, true DMs and RMs in this mouse dataset were obtained by comparing paired RNA-seq samples of wild type (WT) and *Adar1* knockout conditions (Supplementary Fig. 7a, Supplementary Table 9, and Methods), and then used to evaluate the performance of the embedded DeepDDR model in the mouse sample. Since the features used by RED-ML only designed to be extracted from the human genome, RED-ML was failed to be included in this comparison and only EditPredict was applied to the comparison with DeepDDR when analyzing mouse datasets (Fig. 3b). As shown in Fig. 3b, DeepDDR achieved higher AUROC value in the prediction of RMs than EditPredict did; however, the RM recall rate by DeepDDR was only 0.44, when using the same cutoffs as in human (prediction probabilities of DMs ≥ 0.5 and prediction probabilities of RMs < 0.5) (Fig. 3c). Further examination showed that the prediction probabilities of most mouse RMs by DeepDDR were ranged between 0 and 0.8 (Supplementary Fig. 7b), compared to human ones between 0 and 0.5 (Supplementary Fig. 2a and 3a). In this case, the original DeepDDR model trained with human datasets might not be suitable for direct DM and RM classification in mouse datasets.

**Fig. 3.**
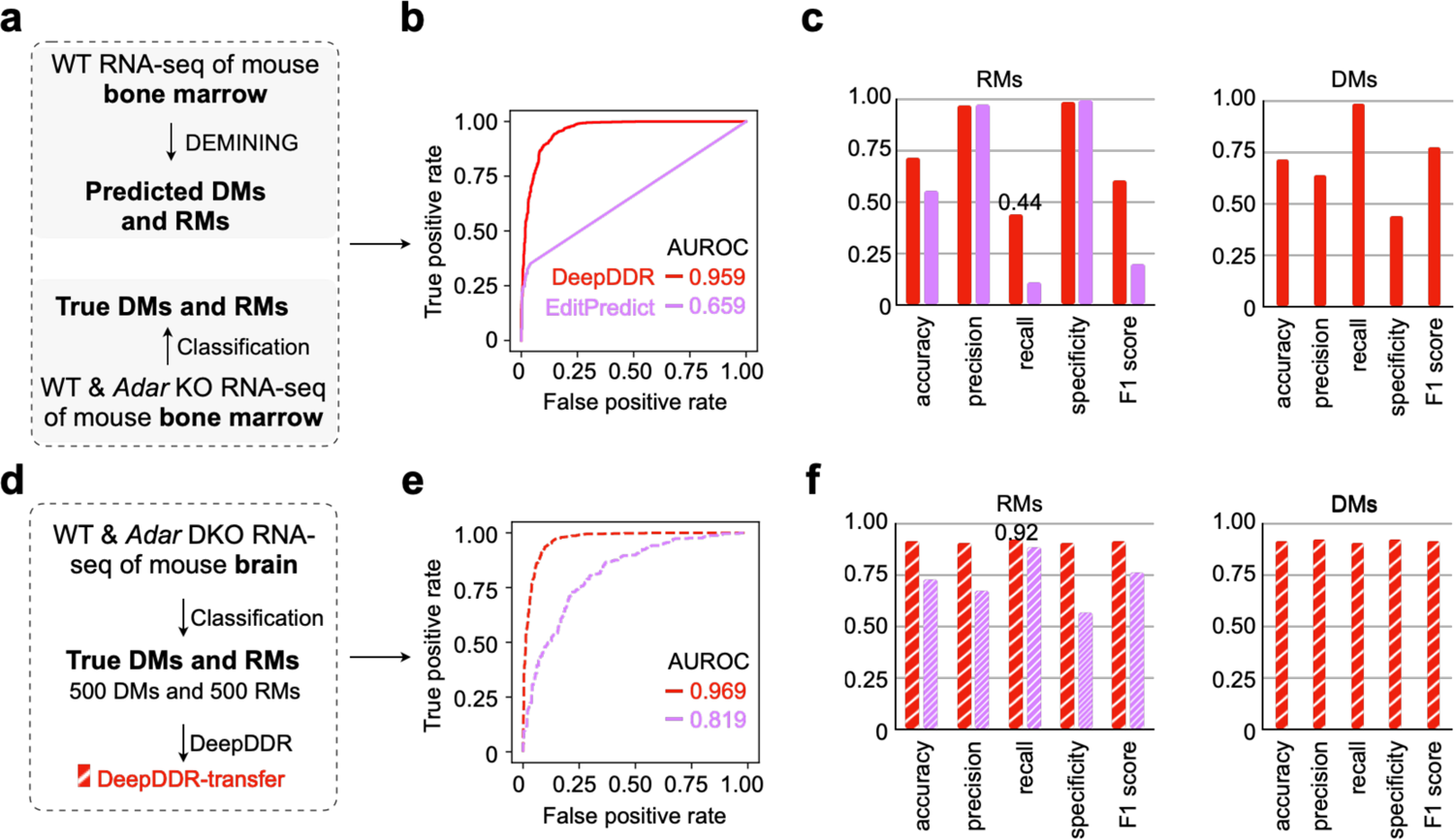
Applying transfer learning to classify DMs and RMs in mouse datasets. **a**, Prediction of DMs and RMs in mouse datasets with original DeepDDR model. DeepDDR was used to predict DMs and RMs from WT RNA-seq data of mouse bone marrow, and true DMs and RMs were classified by comparing WT and *Adar1* KO RNA-seq data. See “Methods” section for details. **b**, Evaluation of different models on RM identification. ROC of DeepDDR (red) and EditPredict (purple) were shown to indicate their performance on RM identification with the mouse bone marrow dataset. AUROC values of DeepDDR (red) and EditPredict (purple) were included in the figure. Of note, since the features used by RED-ML only designed to be extracted from the human genome, RED-ML was failed to be included in this and below comparisons. **c**, Evaluation metrics for RM identification (left) and DM identification (right), including accuracy, precision, recall, specificity, and F1 score, by DeepDDR (red) and EditPredict (purple). **d**, Schematic of constructing DeepDDR-transfer model. An additional mouse brain RNA-seq datasets containing WT and *Adar*s KO samples were used for transfer learning. See “Methods” section for details. **e**, Evaluation of different models after transfer learning on RM identification. ROC of DeepDDR (red dashed line) and EditPredict (purple deashed line) were shown to indicate their performance on RM identification with the same mouse bone marrow dataset. AUROC values of DeepDDR (red) and EditPredict (purple) were included in the figure. **f**, Evaluation metrics for RM identification (left) and DM identification (right), including accuracy, precision, recall, specificity, and F1 score, by DeepDDR-transfer (red shaded bar) and EditPredict-transfer (purple shaded bar).

To solve this problem, we then leveraged the original DeepDDR trained with human datasets as a pre-trained model for transfer learning with another mouse dataset containing both mouse brain *Adar*s (*Adar1* and *Adar2*) knockout and WT samples (Methods). True DMs and RMs in this mouse brain datasets were similarly retrieved by comparing ratio changes between WT and *Adar*s knockout brains samples (Supplementary Fig. 7c, Supplementary Table 10, and Methods), and further used as inputs for the transfer learning to obtain a fine-tuned DeepDDR-transfer model (Fig. 3d and Supplementary Fig. 7d). Similarly, transfer learning was applied to EditPredict using its default requirement of 201-nucleotide sequences extracted around the same true DMs and RMs from mouse brain datasets, resulting in the generation of EditPredict-transfer (Supplementary Fig. 7d). DeepDDR-transfer and EditPredict-transfer were thus used to reanalyze the aforementioned mouse bone marrow WT RNA-seq samples. As shown in Fig. 3e, DeepDDR-transfer exhibited higher AUROC values compared to EditPredict-transfer did. More importantly, the recall ratio of mouse RMs was increased from 0.44 by the original DeepDDR model to 0.92 by the fine-tuned DeepDDR-transfer model (compared Fig. 3f with 3c), indicating that the transfer learning approach significantly improved the model performance. Closer examination showed that prediction probabilities of 90.78% mouse bone marrow A-to-I RMs ranged between 0 to 0.05 by the fine-tuned DeepDDR-transfer model (Supplementary Fig. 7e), compared to those between 0 and 0.8 by the DeepDDR model (Supplementary Fig. 7b).

Similar improvement was also achieved in a downloaded nematode dataset that contains *adr-1* and *adr-2* knockout and WT samples (Methods, Supplementary Table 11), in which the recall ratio was increased from 0.18 by DeepDDR to 0.85 by DeepDDR-transfer (Supplementary Fig. 8). These results together suggested the expanded application of DeepDDR-transfer in the prediction of RNA A-to-I RMs from non-primate samples. Interestingly, when applied the fine-tuned DeepDDR-transfer model back to the human samples, the precision of RM prediction was significantly dropped from 0.94 by the original DeepDDR models (Supplementary Fig. 3d) to 0.56 by DeepDDR-transfer (Supplementary Fig. 9a). Correspondingly, many true DMs from this human dataset were wrongly classified as RMs by DeepDDR-transfer (Supplementary Fig. 9b). In this scenario, the DeepDDR and DeepDDR-transfer models should be individually used for the corresponding analyses in primate or non-primate samples.

During the study, we also developed other machine learning models, such as Light Gradient Boosting Machine (LightGBM), Logistic Regression (LR) and Random Forest (RF), and deep learning models, such as Recurrent Neural Network (RNN) and a hybrid of CNN and RNN (CNN+RNN), for the analyses. Among all these models, DeepDDR with two layers of CNN and the CNN+RNN hybrid model demonstrated comparable performance in distinguishing differentiating DMs and RMs in both human (Supplementary Fig. 10) and mouse datasets, before and after transfer learning (Supplementary Fig. 11-12). Their performance was superior to that of all other models, as shown in Supplementary Fig. 10-12.

## Discussion

Identifying disease-related DMs is of essential to our understanding of genetic disorders contributing to diseases. In the last decade, a large number of WGS and WES datasets have been accumulated to contribute to the profiling of genetic DMs^1, 21^, but suffered with high cost, low efficiency and time-consuming to retrieve high-confidence mutations from increased genomic datasets. Although DMs could be also detected from RNA-seq datasets as DNA transcribes RNA, they were basically treated as noises during RM profiling. Since whole transcriptomic RNA-seq, but not corresponding WGS, datasets are prevalently available from individuals, including patients^22^, we report here the DEMINING framework to directly differentiate expressed DMs and RMs from RNA-seq datasets.

With the embedded DeepDDR model that contains a two-layer CNN architecture, the DEMINING framework showed superior performance over other reported methods like EditPredict^23^ and RED-ML^24^ in the predication of RMs from RNA-seq datasets (Fig. 1). Strikingly, the DEMINING could successfully identify DMs with high efficiency and accuracy when using independent datasets for evaluation (Fig. 1), suggesting its application in the disease-related DM profiling. Importantly, after transfer learning, DEMINING-transfer has exhibited remarkable adaptability in datasets from non-human species, including mouse and nematode (Fig. 3), indicating its broad application in DM and RM profiling across species.

When applied in RNA-seq datasets from AML patient samples^13^, DEMINING uncovered previously unreported DMs and RMs in some uncharacterized AML-associated gene loci, such as *ANKRD36C*, *ANKRD36B* and *ANKRD36B* (Fig. 2). On the one hand, many of recoding DMs are enriched in these *ANKRD* gene regions encoding IDRs, thus suggesting a possible link between dysregulated LLPS and AML pathogenesis (Fig. 2). On the other hand, a positive correlation between DEMINING-identified recoding DMs and up-regulated expression of these *ANKRD* mRNAs has been observed, providing potential targets for AML diagnosis and/or therapeutics (Fig. 2). Of special note, some AML-specific recoding mutations, including two DMs and one RM, were found to be responsible for the production of neoantigens validated by corresponding mass spectrometry data (Supplementary Fig. 6). These findings together demonstrated the successful application of DEMINING in identification of disease-associated mutations, providing additional insights in pathogenesis and therapeutics.

In conclusion, we developed a computational framework, DEMINING, empowered by an embedded DeepDDR model, to differentiate DMs and RMs directly from RNA-seq datasets. DEMINING is characteristic to efficiently and specifically detect DMs from RNA-seq datasets. Its application in AML samples successfully identified previously-underappreciated mutations in unreported AML-associated genes. Since whole transcriptomic RNA-seq datasets are prevalently available from individuals, including patients, we expect that, when applied in a broader spectrum of human disease RNA-seq samples, DEMINING will uncover more disease-related mutations and genes for potential diagnosis and therapeutics.

## Methods

### Collection of published WGS and/or RNA-seq datasets for analyses

A series of WGS and RNA-seq datasets from independent resources were collected and used in this study (Supplementary Table 1). The first data collection consisted of data from 403 donors, which represents the intersection of individuals from the 1000 Genomes Project^1^ phase 1, phase 3, and the Geuvadis consortium^11^. For each donor, data from Whole Genome Sequencing (WGS) and/or Whole Exome Sequencing (WES) were obtained in the form of Variant Call Format (VCF) files.

These VCF files were obtained from both phase 1 (1000 Genomes Project phase 1: ftp://ftp.1000genomes.ebi.ac.uk/vol1/ftp/release/20110521) and phase 3 (1000 Genomes Project phase 3: ftp://ftp.1000genomes.ebi.ac.uk/vol1/ftp/release/20130502) of the 1000 Genomes Project and were merged into a single file if they were from one donor.

Additionally, RNA-seq data in FASTQ file format were obtained from the Geuvadis consortium for the same set of 403 donors. These data were used to construct DNA mutation (DM) and RNA mutation (RM) sets for deep learning model training and evaluation.

The second data collection consisted of an additional donor with SampleID HG00145, who was exclusively included in the 1000 Genomes Project phase 3 and the Geuvadis consortium. The genetic mutations obtained from both WGS and WES for this donor were merged into a single VCF file from the 1000 Genomes Project phase 3. Furthermore, RNA-seq data in FASTQ file format for donor HG00145 were obtained from the Geuvadis consortium. These data were used to construct DM and RM sets, which were used as an independent test set for deep learning model evaluation.

The third data collection consisted of RNA-seq datasets of 19 acute myeloid leukemia (AML) patients (peripheral blood or bone marrow tissue) and 17 normal control samples (medullary thymic or myelocytic precursor cells). These RNA-seq datasets were available from the Gene Expression Omnibus (GEO) with accession number GSE147524 and GSE98310^13^, respectively. These data were used to identify disease-related mutations in AML by DeepDDR-embedded DEMINING framework.

The fourth data collection consisted of mouse bone marrow RNA-seq of three *Adar1* knockout and three wild-type (WT) samples^25^, mouse brain RNA-seq of three *Adar1* and *Adar2* knockout and three WT samples^26^, and nematode RNA-seq of four *adr-1* and *adr-2* knockout and four WT samples^27^. These data were used to construct DeepDDR-transfer model and identify DMs and RMs from other non-primate by DEMINING framework. Detailed information of all these WGS and/or RNA-seq datasets was listed in Supplementary Table 1.

### Identification of true RMs from 403 human RNA-seq datasets by the RADAR pipeline

The previously-published RADAR pipeline^12^ was applied to identify high-confidence RMs from each of 403 RNA-seq datasets, including two major steps.

Step 1: RNA-seq read mapping/piling

After performing quality checked by FastQC (version 0.11.4) with default parameters, RNA-seq reads were trimmed by Trimmomatic (version 0.36) with the following parameters: TruSeq3-PE-2.fa:2:30:10 TRAILING:25 MINLEN:30. Reads mapped to ribosomal DNA (rDNA) by BWA-MEM algorithm (version 0.7.9a) with default parameters were then removed. To enhance the detection of mutations from reads with more mismatches, a two-round unique mapping strategy^10^ was implemented to align high-quality RNA-seq reads to the human hg38 reference genome with the GENCODE gene annotation (version 41) by HISAT2 (version 2.1.0) with parameters: --rna-strandness RF --no-mixed --secondary --no-temp-splicesite --known-splicesite-infile --no-soft-clip --score-min L,-16,0 --mp 7,7 --rfg 0,7 --rdg 0,7 --max-seeds 20 -k 10 --dta and BWA-MEM (version 0.7.9a) with default parameters. Uniquely mapped RNA-seq reads by HISAT2 and BWA-MEM with up to six mismatches were selected and combined for subsequent analysis. The combined reads were analyzed to identify read pairs that were likely derived from duplicates of the same original DNA fragments due to artifactual processes.

As these duplicates were considered non-independent observations, we employed Picard MarkDuplicates (version 2.7.1) with the following parameters: CREATE_INDEX = true and VALIDATION_STRINGENCY = SILENT. This tool tagged all but a single read pair within each set of duplicates, causing the marked pairs to be ignored during subsequent processing steps.

To address issues arising from RNA-seq reads containing Ns in their cigar string, we utilized the GATK (version 4.1.2.0) command SplitNCigarReads with default parameters. This command split reads that contain Ns in their cigar string, typically associated with spanning splicing events in RNA-seq data. The process involved identifying all N cigar elements and generating k+1 new reads, where k represents the number of N cigar elements detected. The first read of generated k+1 new reads included the bases that located to the left of the first N element, while the part of the read that located to the right of the N (including the Ns) was hard clipped and so on for the rest of the new reads.

Base quality scores were crucial for assessing the reliability of base calls and were used to weigh the evidence for or against potential mutations during the mutation discovery process. Systematic biases that affected base quality scores could arise from various factors during the generation of RNA-seq data such as library preparation, sequencing, chip manufacturing defects, or sequencer instrumentation defects. To address systematic biases, we applied a base quality score recalibration procedure using two GATK commands: BaseRecalibrator and ApplyBQSR, with default parameters. During the recalibration process, covariate measurements from all uniquely mapped reads were collected, including read group, base quality score, machine cycle producing the base (Nth cycle = Nth base from the start of the read), and the current base in conjunction with the previous base (dinucleotide). These covariate measurements were used to build a model that captured the systematic biases present in the data. Based on this model, base quality adjustments were applied to the dataset to correct for the observed biases.

Mutation sites were determined from the BAM file containing uniquely mapped reads by using the Samtools (version 1.9) command mpileup with parameters: -Q 0. High-confidence mutations with base quality ≥ 20, mutation reads ≥ 2, hits per billion mapped bases (HPB) ≥ 3, and mutation frequencies ≥ 0.05 were selected for subsequent analyses.

Step 2: Identification of true RMs

To identify high-confidence RMs, expressed mutation sites (with base quality ≥ 20, mutation reads ≥ 2, HPB ≥ 3, and mutation frequencies ≥ 0.05) which were overlapped with DMs listed by 1000 Genomes Project were first removed, and non-overlapped ones were then classified into *Alu* or non-*Alu* regions according to the genomic location as previously described^8^.

Next, mutations in non-*Alu* regions were subjected to additional stringent filtering processes to remove false positives, including:

1. those overlapped with SNPs from dbSNP version 151 (https://www.ncbi.nlm.nih.gov/SNP/), the 1000 Genomes Project (https://www.internationalgenome.org/), or the University of Washington Exome Sequencing Project (https://evs.gs.washington.edu/EVS/)
2. those in simple repeats, homopolymer runs of ≥ 5 base pairs
3. intronic candidates if they were located within 4 base pairs of a known splice junction
4. those mapped to highly similar regions by BLAST-like alignment tools (BLAT^28^, version 364, parameters: -repMatch = 2253 -stepSize = 5)

RMs in non-*Alu* regions after this stringent selection were combined with those in *Alu* regions to obtain high-confidence RMs in each RNA-seq dataset. Of note, over 98% of identified RMs in each of 403 RNA-seq datasets were A-to-G and T-to-C mismatches, mainly in *Alu* regions, indicating high-confidence A-to-I^7^ called by RADAR^12^ (Supplementary Fig. 1b). A total of 122,872 unique A-to-I sites were finally determined from the 403 RNA-seq datasets as the RM set, which is divided into training, validation, and test sets using an 8:1:1 ratio for the development and evaluation of DeepDDR (Supplementary Fig. 1d). Detailed information of these RMs was listed in Supplementary Table 2.

### Identification of true DMs from VCF files of 403 paired human WGS datasets

True DMs were directly obtained from about 31 million SNPs documented in VCF files from the 1000 Genomes Project^1^ (see “Collection of published WGS and/or RNA-seq datasets for analyses” section for detail). Expressed DMs were then determined with base quality ≥ 20, mutation reads ≥ 2, HPB ≥ 3, and mutation frequencies ≥ 0.05 from their corresponding human RNA-seq datasets, as described in “Step 1: RNA-seq read mapping/piling” of the “Identification of true RMs from 403 human RNA-seq datasets by the RADAR pipeline” section. In total, from approximately 578 thousand expressed DMs, a subset of 122,872 expressed DMs was randomly selected and further divided into training, validation, and test sets using an 8:1:1 ratio for the development and evaluation of DeepDDR (Supplementary Fig. 1d). Detailed information of these 122,872 expressed DMs was listed in Supplementary Table 3.

### Construction of a deep learning model for the differentiation of expressed DMs from RMs directly from aligned RNA-seq reads

To differentiate expressed DMs from RMs directly from aligned RNA-seq reads, a deep learning neural network, called DeepDDR, was trained with true DM and RM sets identified from 403 paired human WGS and RNA-seq datasets as described above in “Identification of true DMs from VCF files of 403 paired human WGS datasets” and the “Identification of true RMs from 403 human RNA-seq datasets by the RADAR pipeline” sections, respectively.

To train DeepDDR, the <u>c</u>o-occurrence frequencies of each <u>m</u>utation site with its <u>c</u>ontext bases (CMC) were determined by analyzing 51-nucleotide sequences containing the mutation in the middle and its 50 context bases (25 bases in the upstream region and 25 bases in the downstream region). Briefly, 51-nucleotide sequences with the mutation in the middle were extracted from aligned RNA-seq reads as described in “Step 1: RNA-seq read mapping/piling” of the “Identification of true RMs from 403 human RNA-seq datasets by the RADAR pipeline” section. A 16 x 50 CMC matrix, indicating frequencies of all sixteen types of dinucleotides by 50 combinations of the given DM/RM individually with each of 25 upstream and 25 downstream bases, was obtained from piled 51-bp sequencing, shown in Fig. 1b. CMC matrices of all DMs and RMs in training set were then used as the input sets for the DeepDDR model training.

The DeepDDR model consists of two layers of convolutional neural networks (CNNs) implemented with the TensorFlow v2.0.0 backend in Python v3.7.8. Using CMC matrices as input, the CNN module captures the specific co-occurrence frequencies around DMs or RMs. The model architecture consists of two convolutional layers with Rectified Linear Unit (ReLU) activation function, followed by a max pooling layer. The hyperparameters of the model were optimized through a search process. Specifically, we explored different values for the number of filters in the first convolutional layer, choosing from options such as 32, 64, and 128. Additionally, we varied the number of units in the dense layer, selecting values such as 256, 512, and 1024. During the optimization process, we evaluated the models based on their AUROC (area under the receiver operating characteristic curve) performance on the validation set, as described in “Step 1: RNA-seq read mapping/piling” of the “Identification of true RMs from 403 human RNA-seq datasets by the RADAR pipeline” section. The best-performing model had the following configuration: the first convolutional layer had 128 filters with a width of 12 bases and 4 channels. The second convolutional layer had 64 filters, covering the 128 output channels from the first layer, with filters of width 6 bases. The max pooling layer subsampled the signal with a stride of 4. The resulting signal was then passed to a dense layer with 1024 hidden units, incorporating a dropout ratio of 0.3 to prevent overfitting. The dense layer was connected to two output nodes using the softmax activation function, which allowed for the prediction of probabilities for the two classes: DM or RM. To make a prediction, DeepDDR selected the class with the highest prediction probability as the predicted class. Mutations with prediction probabilities ≥ 0.5 were considered as DMs, mutations with prediction probabilities < 0.5 were considered as RMs.

The DeepDDR model was trained by the CMC matrices of DMs and RMs with a batch size of 5000 and 100 epochs. Early stopping was employed with a patience of 10 rounds, terminating the training process if no improvement in performance was observed. The model with best performance during training was kept at last.

### Construction of the DEMINING framework

DEMINING is a stepwise computational framework to classify DMs and RMs directly from RNA-seq data, including two major steps, RNA-seq read mapping/piling to call high confidence mutations, which are further classified as DMs and RMs by DeepDDR. Briefly, each RNA-seq dataset was first aligned and piled up as described in “Step 1: RNA-seq read mapping/piling” of the “Identification of true RMs from 403 human RNA-seq datasets by the RADAR pipeline” section. After this step, high-confidence mutations were extracted with base quality ≥ 20, mutation reads ≥ 2, HPB ≥ 3, and mutation frequencies ≥ 0.05. Next, these identified high-confidence mutations were then fed into the DeepDDR model for DM and RM classification as described in the “Construction of a deep learning model for the differentiation of expressed DMs from RMs directly from aligned RNA-seq reads” section.

Basically, DeepDDR analyzed the CMC of each input mutation and assigned it a predicted probability score. Mutations with prediction probabilities ≥ 0.5 were considered as DMs, and mutations with prediction probabilities < 0.5 were considered as RMs.

### True DMs and RMs obtained from HG00145 as an independent test set

The same RADAR pipeline^12^ was applied to identify 5,348 RMs from an independent donor with SampleID HG00145. With its paired WGS (see “Collection of published WGS and/or RNA-seq datasets for analyses” section for detail), 17,230 expressed DMs were also retrieved from 1000 Genomes Project and detected in the RNA-seq dataset with base quality ≥ 20, mutation reads ≥ 2, HPB ≥ 3, and mutation frequencies ≥ 0.05. These DMs and RMs were used as independent sets for deep learning model validation and comparison. Detailed information of these 5,348 RMs and 17,230 DMs were listed in Supplementary Table 4.

### Prediction of RMs by EditPredict and RED-ML

For EditPredict, the scripts “get_seq.py” and “editPredict.py” were obtained from the EditPredict GitHub repository (https://github.com/wjd198605/EditPredict). The “get_seq.py” script was used to extract the flanking 200bp sequence of each mutation from the hg38 reference genome. This was done by running the script with the following parameter: “python get_seq.py -f hg38.fa -m b -l 200”. The extracted sequences were then fed to the “editPredict.py” script, which was used to predict RNA editing sites in the input sequences. The EditPredict weights and construction files were obtained from the same repository.

For RED-ML, the RED-ML executable commands “MismatchStat”, “MutDetML” and the script “red_ML.pl” were obtained from the RED-ML GitHub repository (https://github.com/BGIRED/RED-ML). The features used by RED-ML for each mutation were extracted using the “MutDetML” command. This was done with the following parameters: “MismatchStat -i $RNA_BAM -o stat.txt -u -q 20” and “MutDetML -i $RNA_BAM -r hg38.fa -v stat.txt -u -q 20 -Q 0 -d 0 -a 0 -t -o Mut.txt.gz”. The extracted features were then used as input to the “red_ML.pl” script to predict A-to-I editing. Of note, RED-ML was only designed to extract features from the human genome for human A-to-I prediction.

### Gene expression analysis

Gene expression of AML RNA-seq samples was determined by StringTie (version 2.1.4) with default parameters on BAM files after RNA-seq read mapping, as previously described^29^.

### Gene ontology analysis

After annotating mutations using the ANNOVAR^30^ (version 7 Jun 2020), GO analyses of three gene sets, including all 4,464 genes containing 8,220 recoding DMs, 86 AML-associated genes listed in CGC, and their overlapped 50 genes, were performed by using clusterProfiler enrichGO function (ont = “BP”, pAdjustMethod = “BH”, pvalueCutoff = 0.2, qvalueCutoff = 0.2) and reduced redundancy of GO terms by simplify function (cutoff = 0.7, by = “p.adjust”, select_fun = min, measure = “Wang”). Top GO terms ordered by adjusted P value in each gene set were kept and compared (Fig. 2e).

### Mass spectrometric data analysis

Raw MS data of major histocompatibility complex (MHC) associated peptides from 19 AML patient were available in a public repository from the ProteomeXchange Consortium via the PRIDE partner repository by the dataset identifier PXD018542^13^. To analyze these raw MS data, we constructed an *in silico* library containing predicted peptides ranging from 8 to 30 amino acids in length from 4,464 genes with recoding DMs and 121 genes with recoding RMs. Decoyed peptides from the predicted peptides and proteins from human reference proteome (UniProt 2023-04-04) were also added to this library to be used as negative controls. Subsequently, the *in silico* library was served as a reference database for searching LC-MS/MS spectra. Raw MS data of each sample was transformed into mzML format using MSConvert^31^ (version 3.0.23090). The mzML format of MS data was then searched against the customized database using Comet^32^ (version 201901 rev. 1). The search was performed with 10 ppm precursor mass tolerance and 0.02 fragment ions mass tolerance. The search results were analyzed by PeptideProphet^33^ (version 5.2.1). To ensure a reliable peptide identification, the search results were filtered to achieve an estimated peptide FDR of 5%.

Only mutation-containing peptides that did not occur in the reference proteome were included in the final list. The search and filtering processes were implemented by the Philosopher (version 4.8.1) framework^34^. The detailed information of the search results was listed in Supplementary Fig. 6b.

### Identification of true DMs and RMs from paired adenosine deaminase(s) knockout and wild-type samples

To identify true DMs and RMs from adenosine deaminase(s) knockout (KO) and wild-type (WT) data, the following steps were performed:

Step 1: RNA-seq data from the adenosine deaminase(s) KO and WT samples were mapped to the corresponding reference genomes (mouse: mm10, nematode: ce11) using the same method described in “Step 1: RNA-seq read mapping/piling” of the “Identification of true RMs from 403 human RNA-seq datasets by the RADAR pipeline” section.

Step 2: Mutations were determined from the BAM file containing uniquely mapped reads by using the Samtools (version 1.9) command mpileup with parameters: -Q 0. High-confidence mutations with base quality ≥ 20, mutation reads ≥ 2, HPB ≥ 3, and mutation frequencies ≥ 0.05 were selected for further analyses.

Step 3: Calculating mutation frequency ratio

Firstly, calculate the average mutation frequency in the wild-type (WT) samples by summing up the mutation frequencies of all the WT samples and dividing by the total number of WT samples. Secondly, calculate the average mutation frequency in the adenosine deaminase(s) knockout (KO) samples by summing up the mutation frequencies of all the KO samples and dividing by the total number of KO samples. Thirdly, subtract the average mutation frequency of the WT samples from the average mutation frequency of the KO samples.

Mutation frequency difference = Average mutation frequency in KO samples - Average mutation frequency in WT samples Lastly, divide the difference obtained in last step by the average mutation frequency in the WT samples. Mutation frequency ratio = Mutation frequency difference / Average mutation frequency in WT samples

Step 4: Classification of true DMs and RMs:

Ture DMs were selected by criteria: 1) the absolute value of mutation frequency ratio < 0.2, 2) the mutation frequencies in WT samples ≥ 0.05, and 3) not overlapped with sites in the REDIportal database^35^.

Of note, as the REDIportal did not contain nematodes, ture DMs from nematode dataset^27^ were only selected by the absolute value of mutation frequency ratio < 0.2 and the mutation frequencies in WT samples ≥ 0.05.

Ture RMs were selected by criteria: 1) The mutation frequency ratio were lower than −0.2; 2) The mutation frequencies in WT samples were greater than 0.05 and lower than 0.5; 3) The mutation frequencies in adenosine deaminase(s) KO samples were lower than 0.05; 4) The mutation types were A-to-G or T-to-C, indicating an A-to-I editing event; 5) The mutations did not overlap with SNPs in the dbSNP database (dbSNP version for mouse: 150, dbSNP version for nematode: 138). Detailed information of these RMs and DMs were listed in Supplementary Table 9-11.

### Fine-tuning of DeepDDR by transfer learning

To perform fine-tuning and validation of DeepDDR on mouse bone marrow datasets using transfer learning, the following steps were taken:

Step 1: Obtain true DMs and RMs from the mouse bone marrow datasets using the method described in the “Identification of true DMs and RMs from paired adenosine deaminase(s) knockout and wild-type samples” section.

Step 2: Split the DMs and RMs individually into transfer training, transfer validation, and transfer test sets with a ratio of 70:15:15.

Step 3: Sequentially select transfer training sets of different sizes from the original 70% split transfer training set. The aim is to find the smallest transfer training set size that maximizes the recall ratio. Evaluate the recall ratio and other evaluation metrics on the transfer validation sets to determine the optimal size.

Step 4: Based on the evaluation results, select the subset of 500 DMs and 500 RMs as the final transfer training set.

Step 5: Construct the DeepDDR-transfer model using a transfer learning strategy. Start with the DeepDDR model trained on human datasets as described in the “Construction of a deep learning model for the differentiation of expressed DMs from RMs directly from aligned RNA-seq reads” section. Fine-tune this model using the final transfer training set from the mouse bone marrow datasets. This process was implemented with the TensorFlow v2.0.0 backend in Python v3.7.8.

Step 6: Test the performance of the DeepDDR-transfer model on the transfer test set.

### Construction of other deep learning models

To train other deep learning-based models for comparison, a Recurrent Neural Network (RNN) and a hybrid model of Convolutional Neural Network and RNN (CNN+RNN) were constructed using the TensorFlow (version 2.0.0) backend in Python (version 3.7.8).

For the RNN model, it consisted of one Bidirectional LSTM layer and one dense layer. The hyperparameters that were varied during training include the number of hidden units in the Bidirectional LSTM layer, such as 16, 32, or 64, and the number of units in the dense layer, such as 256, 512, or 1024.

For the CNN+RNN model, it combined two convolutional layers, one Bidirectional LSTM layer, and one dense layer. The hyperparameters that were varied include the number of filters in the first convolutional layer, such as 32, 64, or 128, and the number of hidden units in the Bidirectional LSTM layer, such as 16, 32, or 64.

During training, the models were fed with the CMC matrix as input. The performance of each model was evaluated based on the AUROC (area under the receiver operating characteristic curve) on the validation set. The model that achieved the highest AUROC on the validation set was chosen for further comparison.

### Construction of machine learning models

To train the machine learning models, including Light Gradient Boosting Machine (LightGBM), Logistic Regression (LR), and Random Forest (RF), four features were extracted from the mutations in the training set, as described in “Step 1: RNA-seq read mapping/piling” of the “Identification of true RMs from 403 human RNA-seq datasets by the RADAR pipeline” section. These features include mutation frequency, base substitution type, whether the mutation is located in the *Alu* region, and the count of base pairs from the closest mutation in the genomic sequence. These features were selected based on insights obtained from previous bioinformatic tools and the current understanding of RNA editing mechanisms.

For the LightGBM model, the LightGBM Python package (version 3.1.1.99) was used. For LR and RF models, the Python scikit-learn package (version 0.20.3) was employed. The extracted four features were used as inputs for these models.

To optimize the hyperparameters of each model, 150 combinations were selected for LightGBM, Logistic Regression (LR), and Random Forest (RF) models, individually. For LightGBM, we searched 150 combinations chosen from following hyperparameter configurations: learning rate (chosen from [0.1, 0.05, 0.02, 0.01]), the number of estimators (chosen from 24 points that were evenly spaced between 100 and 2400), the maximum depth of the individual estimators (chosen from [2–5, 10, 20, 40, 51]), the minimum number of data in one leaf (chosen from 22 points that were evenly spaced between 1 and 44), the fraction of subset on each estimator (chosen from [0.2, 0.3, 0.4, 0.5, 0.6, 0.7, 0.8, 0.9]), the frequency of bagging (chosen from [0, 1, 2]), the penalty of L1 regularization (chosen from [0, 0.001, 0.005, 0.01, 0.1]) and the penalty of L2 regularization (chosen from [0, 0.001, 0.005, 0.01, 0.1]). For LR, the penalty of L2 regularization was randomly chosen from integer between 50 and 1000 to optimize hyperparameters. For RF, we searched 150 combinations through the following hyperparameter configurations: the number of trees in the forest (randomly chosen from integer between 1 and 250) and the maximum depth of the tree (randomly chosen from integer between 5 and 255).

To select the optimal hyperparameters for each model, a five-fold cross-validation based on RandomizedSearchCV was performed. The F1 score was used as the evaluation metric, and the model that achieved the highest mean F1 score during cross-validation was chosen for further comparison and analysis.

## Statistical analyses

Statistically significant differences were assessed as described in correspondent figure legends. All statistic tests were performed with the R platform (version 4.1.1) (http://www.R-project.org/).

## Data availability

The statements of data availability and their associated accession codes and references, are available in the “Methods” section and Supplementary Table 1. Multiple datasets from independent resources were used in this study. The first collection consisted of WGS/WES and RNA-seq datasets of 403 donors was available from 1000 Genomes Project phase 1, phase 3, and the Geuvadis consortium^11^. The second collection consisted of WGS/WES and RNA-seq datasets of an additional donor was available from 1000 Genomes Project phase 3 and the Geuvadis consortium. The third data collection consisted of RNA-seq datasets of 19 acute myeloid leukemia (AML) patients (peripheral blood or bone marrow tissue) and 17 normal control samples (medullary thymic or myelocytic precursor cells) was available from the Gene Expression Omnibus (GEO) with accession number GSE147524 and GSE98310^13^. The raw mass spectrometric dataset of major histocompatibility complex (MHC) associated peptides from 19 AML patient was available from ProteomeXchange Consortium with accession number PXD018542^13^. The final collection consisted of RNA-seq of mouse bone marrow, mouse brain RNA-seq and nematode was available from the Sequence Read Archive with project ID PRJEB31568^25^, PRJNA546532^26^ and PRJNA215361^27^, respectively.

## Code availability

All scripts used in this study, including DEMINING framework, DeepDDR model, and related codes are temporarily available at https://github.com/fzc1997/DEMINING/. All these framework, model and codes will be publicly available in the future. Further information and requests for resources should be directed to and will be fulfilled by the corresponding author, Li Yang.

## Acknowledgements

We thank Yang laboratory for discussion. This work was supported by National Natural Science Foundation of China (NSFC, 31925011), the Ministry of Science and Technology of China (MoST, 2019YFA0802804, 2021YFA1300503) and the Chinese Academy of Sciences (CAS, XDB38040300) to L.Y., and by China National Postdoctoral Program for Innovative Talents (BX20220077) and Shanghai Post-doctoral Excellence Program, China (2022728) to F.N.

## Author contributions

L.Y. conceived and supervised the project; Z.-C.F. and B.-Q.G. preformed most analyses with the help of F.N. and X.-K.M., supervised by L.Y. and L.Y. wrote the paper with input from all authors. All authors read and approved the final manuscript.

