## Supplementary Material for "A deep learning model embedded framework to distinguish DNA and RNA mutations directly from RNA-seq"

### SUPPLEMENTARY FIGURES

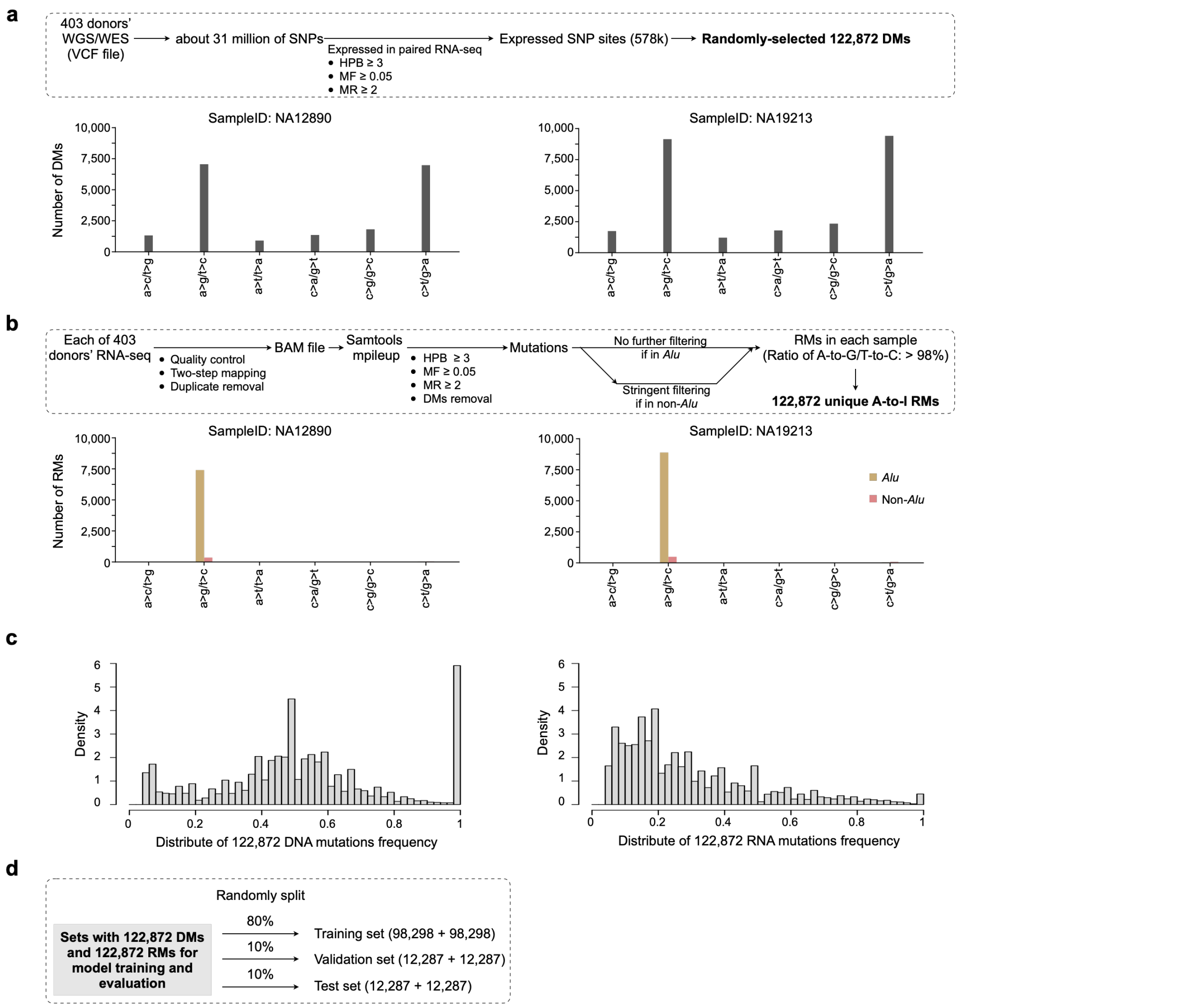


**Supplementary Fig. 1 | Construction of high-confidence DNA and RNA mutations for model development and evaluation.**

**a**, The DNA mutations selection for DeepDDR developing. Top, schematic of a stepwise pipeline for identifying DMs. Bottom, histogram to show numbers of all 6 types of DMs, exemplified by samples NA12890 (left) and NA19213 (right) samples. See "Methods" section for details.

**b**, The RNA mutations selection for DeepDDR developing. Top, schematic of a stepwise pipeline for identifying RMs. Bottom, histogram to show numbers of all 6 types of RMs found in different genomic locations, exemplified by sample NA12890 (left) and NA19213 (right) samples. See “Methods” section for details.

**c**, Distribution of DMs and RMs from 403 donors. Left, mutation frequency distribution for all 122,872 DMs identified in **a**. Right, mutation frequency distribution for all 122,872 RMs identified in **b**.

**d**, Generation of training, validation, and test sets. To facilitate the development and evaluation of DeepDDR, the set of 122,872 DMs and 122,872 RMs were individually split into training, validation, and test sets using an 8:1:1 ratio.

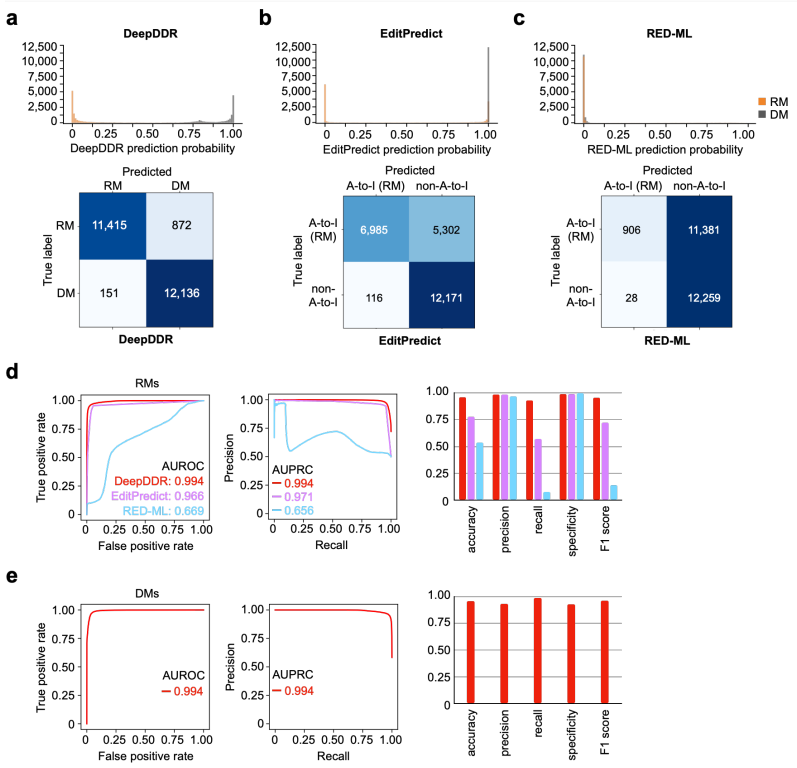


**Supplementary Fig. 2 | DeepDDR shows better performance than other models on test set.**

**a-c**, Distribution of prediction probability (top) and confusion matrix (bottom) for DeepDDR (**a**), EditPredict (**b**) and RED-ML (**c**).

**d**, Evaluation of different models on RM identification. Left, Receiver Operating Characteristic (ROC) curves for RM identification comparing DeepDDR (red), EditPredict (purple) and RED-ML (blue). Area Under ROC (AUROC) values of DeepDDR (red), EditPredict (purple) and RED-ML (blue) were included in the figure. Middle, Precision-Recall Curves (PRC) for RM identification comparing DeepDDR (red), EditPredict (purple) and RED-ML (blue). Area Under PRC (AUPRC) values of DeepDDR (red), EditPredict (purple) and RED-ML (blue) were included in the figure.

Right, evaluation metrics including accuracy, precision, recall, specificity and F1 score for DeepDDR (red), EditPredict (purple) and RED-ML (blue).

**e**, Evaluation of DeepDDR on DM identification. Left, ROC curve for DM identification of DeepDDR. AUROC value of DeepDDR was included in the figure. Middle, PRC for DM identification of DeepDDR. AUPRC value of DeepDDR was included in the figure. Right, evaluation metrics including accuracy, precision, recall, specificity and F1 score for DeepDDR. Of note, since EditPredict and RED-ML could not identify DM, the ROC and PRC for DM identification of EditPredict and RED-ML could not be calculated.

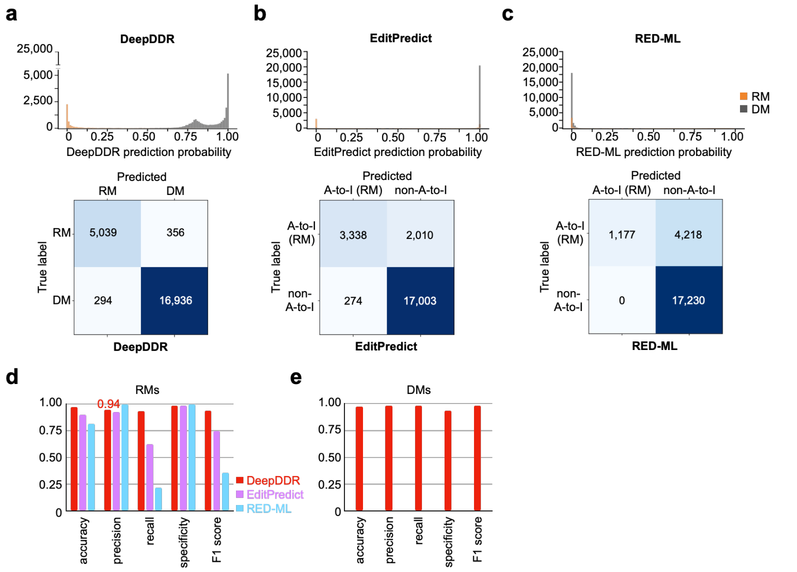


**Supplementary Fig. 3 | DeepDDR shows better performance than other models on independent sample HG00145.**

**a-c**, Distribution of DMs and RMs prediction probability (top) and confusion matrix (bottom) from DeepDDR (**a**), EditPredict (**b**) and RED-ML (**c**).

**d**, Evaluation metrics for RM identification, including accuracy, precision, recall, specificity and F1 score, comparing DeepDDR (red bar), EditPredict (purple bar) and RED-ML (blue bar).

**e**, Evaluation metrics for DM identification, including accuracy, precision, recall, specificity and F1 score, for DeepDDR.

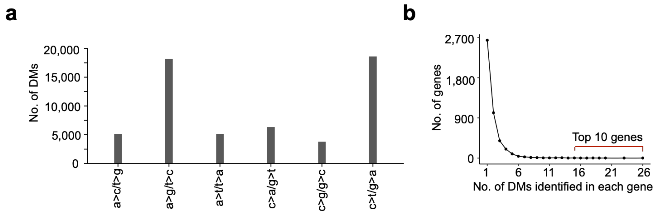


**Supplementary Fig. 4 | Mutation type of AML-specific DMs and number of recoding DMs in mutated genes.**

**a**, Histogram to show the number of all 6 types of AML-specific DMs.

**b**, Distribution of AML-specific recoding DM number in 4,464 mutated genes. The top 10 genes contain more than 15 DMs, labeled with red line.

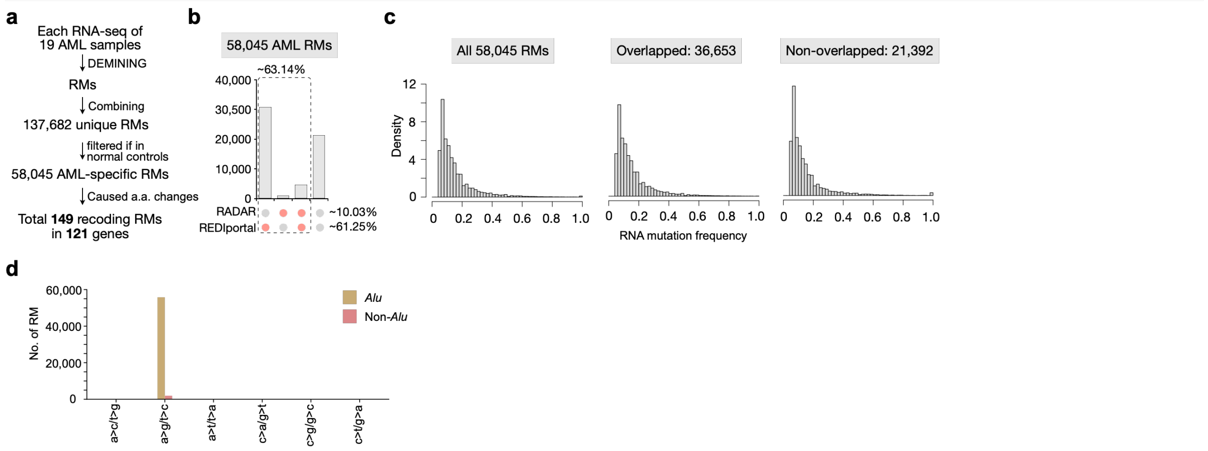


**Supplementary Fig. 5 | Applying DEMINING framework to identify AML-specific RMs.**

**a**, Identification of AML-associated DMs from corresponding RNA-seq datasets by DEMINING.

**b**, Overlapping of the AML-specific RMs with reported RNA editing sites in public databases, including RADAR (http://RNAedit.com) and REDIportal (http://srv00.recas.ba.infn.it/atlas/).

**c**, Mutation frequency distribution of all AML-specific RMs (left), overlapped (middle) and non-overlapped (right) AML-specific RMs.

**d**, Histogram to show the number of all 6 types of AML-specific RMs in different genomic locations.

**a**

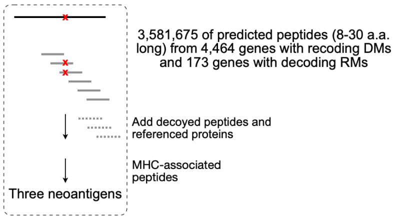


**b**

| sample_id | peptide | length (a.a.) | a.a. mutation position | a.a. mutation | e_value | posibillity | mutation site | mutation frequency | HPB | gene | mutation type | reported database |
|---|---|---|---|---|---|---|---|---|---|---|---|---|
| 05H143 | TSQI**R**AILL | 9 | 5 | p.Q317R | 0.0022 | 0.964 | chr11:5664119 (a>g) | 0.0506 | 7.35 | TRIM5 | RM | REDIportal |
| 14H124 | LTGPGVAGYL TAGTSTSV**I**S NLPPPVDH | 28 | 19 | p.M2225I | 0.043 | 1 | chr3:132538225 (g>a) | 0.4888 | 16.29 | DNAJC13 | DM | dbSNP/ COSMIC/ ClinVar |
| 16H145 | C**H**VYDMRPE TDPQSMFGM RDPPMFIPTP | 28 | 2 | p.Q1586H | 0.0151 | 0.078 | chr7:135626326 (a>c) | 0.56 | 6.31 | NUP205 | DM | dbSNP |

**Supplementary Fig. 6 | Identification of mutated MHC-presented peptides by mass spectrometry (MS) data.**

**a**, Pipeline of identifying mutated MHC-presented peptides. Raw MS data were retrieved from a published study[1].

**b**, Information of neoantigens generated form two DMs and one RM identified by DEMINING.

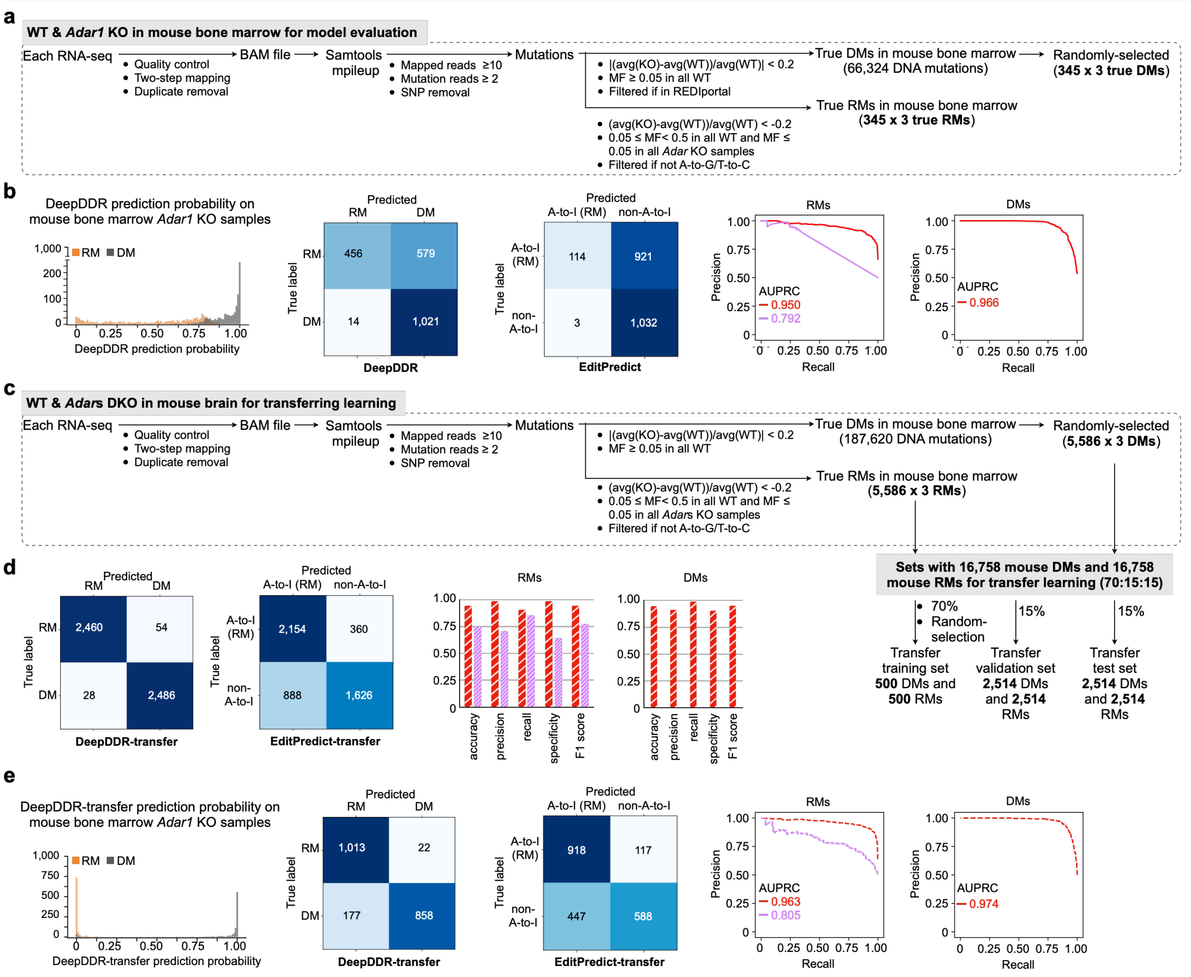


**Supplementary Fig. 7 | Transfer learning and evaluation with mouse datasets.**

**a**, Schematic of a stepwise pipeline used to identify DMs and RMs from the mouse bone marrow test set. See "Methods" section for details.

**b**, Performance of DeepDDR and EditPredict on the mouse bone marrow test set. From left to right: Distribution of prediction probabilities for DeepDDR, confusion matrix for DeepDDR, confusion matrix for EditPredict, PRC for RM identification comparing DeepDDR (red line) and EditPredict (purple line), and PRC for DM identification of DeepDDR.

**c**, Schematic of a stepwise pipeline used to identify DMs and RMs from the mouse brain dataset. The identified DMs and RMs were subsequently divided into transfer training, transfer validation, and transfer test sets using a 70:15:15 ratio. Within the split of the transfer training set, a subset of 500 DMs and 500 RMs were randomly selected based on the findings of a pre-experiment (data not shown). This selection was made as recall began to maximize at approximately 500 DMs and 500 RMs after random subsets within the transfer training set. See "Methods" section for details.

**d**, Performance of DeepDDR-transfer and EditPredict-transfer on transfer test set. From left to right: Confusion matrix for DeepDDR-transfer (red shaded bar) and EditPredict-transfer (purple shaded bar), evaluation metrics for RM identification and DM identification, including accuracy, precision, recall, specificity and F1 score.

**e**, Performance of DeepDDR-transfer and EditPredict-transfer on the same mouse bone marrow test set. From left to right: Distribution of prediction probability for DeepDDR-transfer, confusion matrix for DeepDDR-transfer, confusion matrix for EditPredict-transfer, PRC for RM identification comparing DeepDDR-transfer (red dashed line) and EditPredict-transfer (purple dashed line), and PRC for DM identification of DeepDDR-transfer.

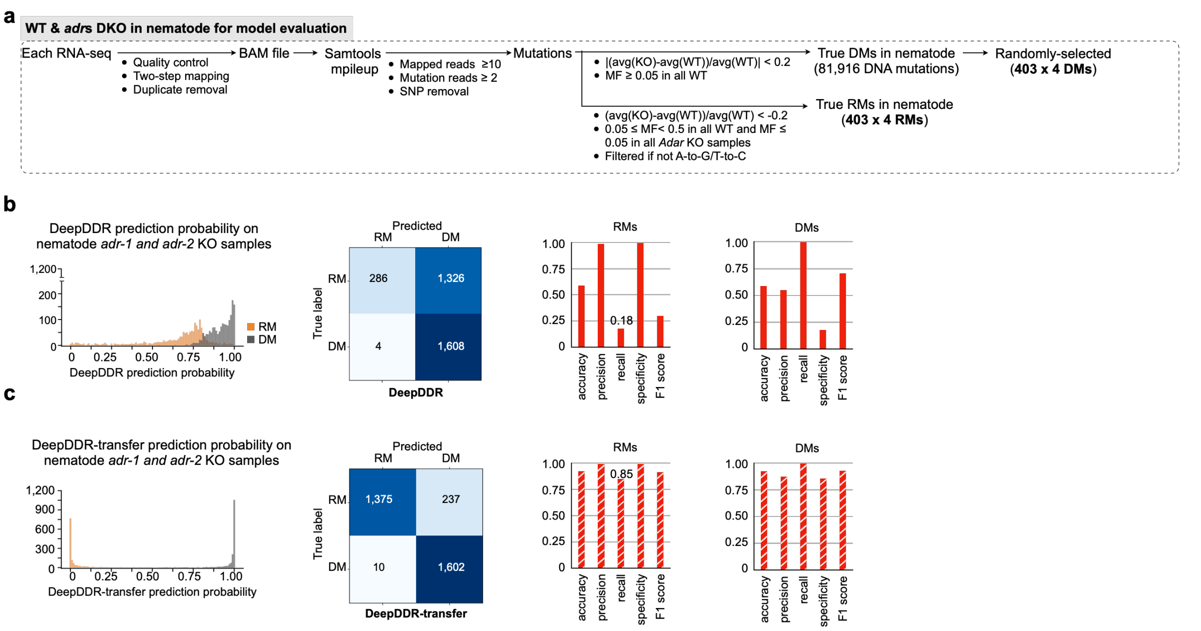


#### Supplementary Fig. 8 | Nematode datasets used for transfer learning and evaluation.

**a**, Schematic of a stepwise pipeline used to identify DMs and RMs from the nematode test set. See “Methods” section for details.

**b**, Performance of DeepDDR on the nematode test set. From left to right: Distribution of prediction probabilities for DeepDDR, confusion matrix for DeepDDR, evaluation metrics for RM identification of DeepDDR and evaluation metrics for DM identification of DeepDDR. The recall of RM identification by DeepDDR was 0.18.

**c**, Performance of DeepDDR-transfer on the nematode test set. From left to right: distribution of prediction probabilities for DeepDDR-transfer, confusion matrix for DeepDDR-transfer, evaluation metrics for RM identification of DeepDDR-transfer and evaluation metrics for DM identification of DeepDDR-transfer. The recall of RM identification by DeepDDR-transfer was 0.85.

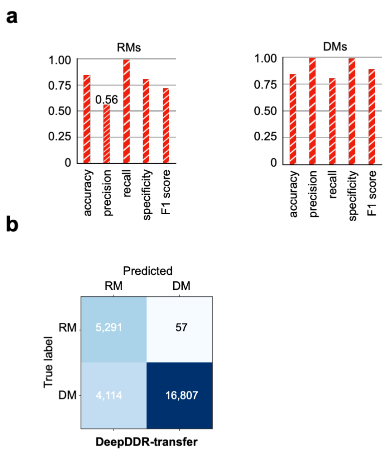


**Supplementary Fig. 9 | Performance of DeepDDR-transfer on HG00145 test set.**

**a**, Performance of DeepDDR-transfer on HG00145 test set. Evaluation metrics for RM identification (left) and DM identification (right), including accuracy, precision, recall, specificity and F1 score, for DeepDDR-transfer. The precision of RM identification by DeepDDR-transfer was 0.56.

**b**, Confusion matrix for DeepDDR-transfer.

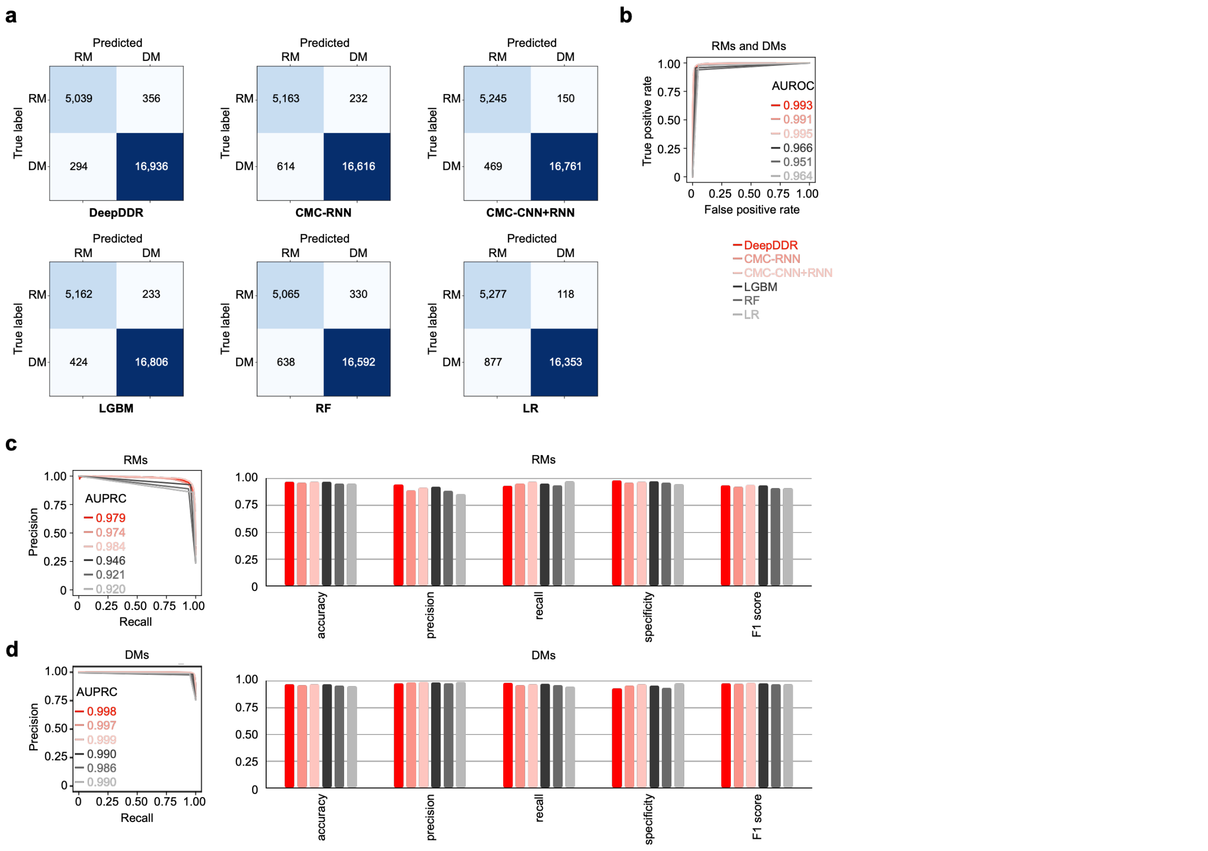


**Supplementary Fig. 10 | Comparison of model architectures on HG00145.**

**a**, Confusion matrix for different model architectures.

**b**, AUROC comparing different model architectures.

**c**, Model performances on RM identification. Left, AUPRC comparing model architectures. Right, evaluation metrics for RM identification, including accuracy, precision, recall, specificity and F1 score.

**d**, Model performances on DM identification. Left, AUPRC comparing model architectures. Right, evaluation metrics for DM identification, including accuracy, precision, recall, specificity and F1 score.

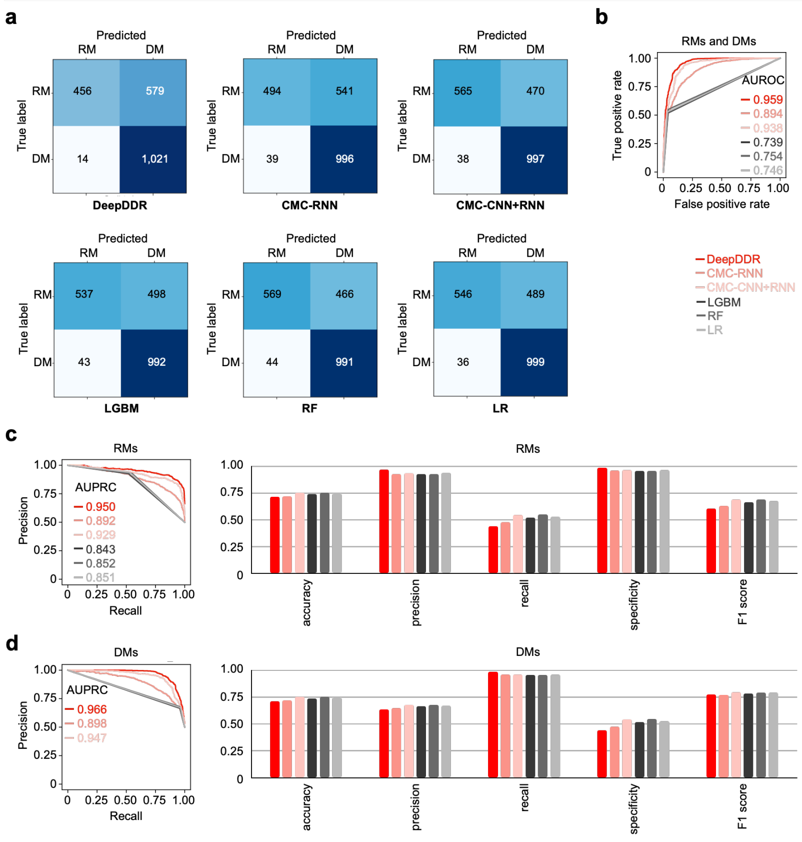


**Supplementary Fig. 11 | Comparison of model architectures on mouse bone marrow test set.**

**a**, Confusion matrix for different model architectures.

**b**, AUROC comparing different model architectures.

**c**, Model performances on RM identification. Left, AUPRC comparing model architectures. Right, evaluation metrics for RM identification, including accuracy, precision, recall, specificity and F1 score.

**d**, Model performances on DM identification. Left, AUPRC comparing model architectures. Right, evaluation metrics for DM identification, including accuracy, precision, recall, specificity and F1 score.

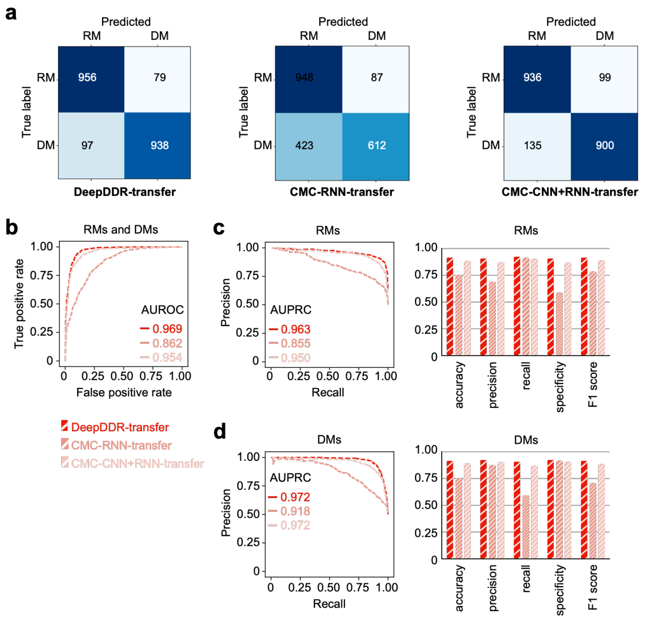


**Supplementary Fig. 12 | Comparison of model architectures on mouse bone marrow test set after transfer.**

**a**, Confusion matrix for different model architectures.

**b**, AUROC comparing different model architectures.

**c**, Model performances on RM identification. Left, AUPRC comparing model architectures. Right, evaluation metrics for RM identification, including accuracy, precision, recall, specificity and F1 score.

**d**, Model performances on DM identification. Left, AUPRC comparing model architectures. Right, evaluation metrics for DM identification, including accuracy, precision, recall, specificity and F1 score.

**SUPPLEMENTARY TABLE LEGENDS**

Supplementary Table 1: Summary of downloaded datasets used in this study.

Include 403 donors' datasets for DeepDDR model establish, two independent validation datasets, RNA-seq data from 19 AML patients and 17 healthy donors and datasets for DeepDDR-transfer training.

Supplementary Table 2: Information of RNA mutations for DeepDDR development and evaluation.

In total, 122,872 RNA mutations (RMs) were used to development and evaluation of DeepDDR. Genomic coordinates (hg38), SampleID, Mutation type, Mutation frequency, Hits per billion mapped bases (HPB), Host gene, Repetitive element, Gene body, Dataset type and Label are included.

Supplementary Table 3: Information of DNA mutations for DeepDDR development and evaluation.

In total, 122,872 DNA mutations (DMs) were used to development and evaluation of DeepDDR. Genomic coordinates (hg38), SampleID, Mutation type, Mutation frequency, Hits per billion mapped bases (HPB), Host gene, Repetitive element, Gene body, Dataset type and Label are included.

Supplementary Table 4: Information of true RMs and DMs from independent HG00145.

In total, 5,395 RMs and 17,230 DMs were identified from HG00145 for deep learning model evaluation. Genomic coordinates (hg38), SampleID, Mutation type, Mutation frequency, Hits per billion mapped bases (HPB), Host gene, Repetitive element, Gene body and Label are included.

Supplementary Table 5: Information of DMs identified in AML patients by DEMINING.

In total, 195,256 DMs were identified by DEMINING in RNA-seq from 19 AML patients. Genomic coordinates (hg38), SampleID, Mutation type, Mutation frequency, Hits per billion mapped bases (HPB) and Condition of AML-specific are included.

Supplementary Table 6: Information of RMs identified in AML patients by DEMINING.

In total, 137,682 RMs identified by DEMINING in RNA-seq from 19 AML patients. Genomic coordinates (hg38), SampleID, Mutation type, Mutation frequency, Hits per billion mapped bases (HPB) and Condition of AML-specific are included.

Supplementary Table 7: Information of AML-specific DMs.

In total, 57,150 AML-specific DMs identified by filtering those also detected in 17 normal control samples or had a high minor allele frequency (MAF > 0.05) in 1000 Genomes Project. Genomic coordinates (hg38), SampleID, Mutation type, Mutation frequency, Hits per billion mapped bases (HPB), Repetitive element, Host gene, Gene body, Condition of recoding, Amino acid change type, Condition of overlapped with ClinVar, COSMIC or dbSNP database are included.

Supplementary Table 8: Information of AML-specific RMs.

In total, 58,045 AML-specific RMs were identified in AML patients by filtering those also detected in 17 normal control samples. Genomic coordinates (hg38), SampleID, Mutation type, Mutation frequency, Hits per billion mapped bases (HPB), Repetitive element, Host gene, Gene body, Condition of recoding, Amino acid change type, Condition of overlapped with RADAR database[2] and Condition of overlapped with REDIportal database[3] are included.

Supplementary Table 9: Information of true RMs and DMs from mouse bone marrow dataset including three *Adar1* knockout and three wild-type samples.

In total, 345 RMs and 345 DMs were identified from mouse bone marrow dataset used to deep learning model evaluation. Genomic coordinates (mm10) and Label are included.

Supplementary Table 10: Information of true RMs and DMs from mouse brain dataset including three *Adar1* and *Andr2* knockout and three wild-type samples.

In total, 5,586 RMs and 5,586 DMs identified from mouse brain dataset used to transfer learning. Genomic coordinates (mm10) and Label are included.

Supplementary Table 11: Information of true RMs and DMs from nematode dataset including four *adr-1* and *adr-2* knockout and four wild-type samples.

In total, 403 RMs and 403 DMs identified from nematode dataset used to deep learning model evaluation. Genomic coordinates (ce11) and Label are included.
